# Uncovering pre-sensitizing agents to FLT3 inhibitors in acute myeloid leukemia with ReSisTrace lineage tracing

**DOI:** 10.1101/2024.10.22.619614

**Authors:** Johanna Eriksson, Shuyu Zheng, Jie Bao, Jun Dai, Wenyu Wang, Anna Vähärautio, Jing Tang

## Abstract

While FLT3 inhibitors have significantly improved the treatment of aggressive FLT3-mutated acute myeloid leukemia (AML), the emergence of resistance remains as a major challenge. Here, we applied our recently developed single-cell lineage-tracing method ReSisTrace to identify cells that are pre-resistant or pre-sensitive to FLT3 inhibitors midostaurin and quizartinib in FLT3-ITD-positive AML. By comparing the gene expression profiles of these cells, we unraveled the transcriptional pre-resistance signatures, including G1 to S phase transition 1 (GSPT1) gene. Targeting GSPT1 with the small molecule CC-90009 exhibited strong synergistic effect when combined with FLT3 inhibitors in the FLT3-ITD-mutated MOLM-13 and MV4-11 cell lines and primary AML patient samples. Further, we identified novel compounds that induced transcriptomic changes opposite to the pre-resistance signatures, thereby driving cells to FLT3 inhibitor-sensitive states. Vistusertib (mTOR inhibitor), linsitinib (IGF1R and insulin receptor inhibitor), and meisoindigo (IGF1R and Src family kinase inhibitor), all inhibiting pathways parallel to or downstream of oncogenic FLT3 signaling, were predicted and validated to pre-sensitize the FLT3-ITD-mutated cell lines and primary cells to FLT3 inhibitors. Collectively, these findings demonstrate the validity of our lineage-tracing method in unveiling pre-existing transcriptional features of treatment vulnerability in hematological cancers, and elucidate novel strategies for enhancing FLT3 inhibitor treatment efficacy in FLT3-ITD-positive AML by preventing the emergence of treatment resistance.

## Introduction

Acute myeloid leukemia (AML) is a highly aggressive and heterogeneous hematological malignancy marked by the infiltration of blood, bone marrow, and tissues by proliferative and abnormally differentiated cells of the myeloid lineage (Döhner et al. 2015). The FMS-like tyrosine kinase-3 (FLT3) is a type 3 receptor tyrosine kinase crucial for the expansion of normal hematopoietic stem cells and it plays a pivotal role in the proliferation and anti-apoptosis of most primary AML cells. FLT3 is activated by binding of the FLT3 ligand (FL), which results in a signaling cascade activating multiple downstream signaling pathways, including PI3K/protein kinase B (AKT) and mitogen-activated protein kinase (MAPK) pathways (Kazi and Rönnstrand 2019). FLT3 is also one of the most commonly mutated genes in AML, with activating internal tandem duplications (ITD) or point mutations in the tyrosine kinase domain (TKD) occurring in approximately 20% and 7% of AML patients, respectively (Yamamoto et al. 2001, Schnittger et al. 2002). In contrast to wild-type FLT3 signaling, FLT3-ITD not only activates PI3K and MAPK signaling but also triggers STAT5 signaling (Kazi and Rönnstrand 2019). The presence of FLT3-ITD mutations also serves as a prognostic marker, indicating a shorter disease-free and overall survival (Sheikhha et al. 2003).

Development of FLT3 inhibitors has changed the standard of care for FLT3-mutated AML. Midostaurin (Levis 2017) and the recently approved quizartinib are the only FDA-approved FLT3 inhibitors as a first-line treatment in combination with chemotherapy for adult patients with newly-diagnosed FLT3-ITD-positive AML. Notably, quizartinib was also approved as maintenance monotherapy for up to 3 years in patients who do not undergo subsequent hematopoietic stem cell transplantation. In addition, gilterinib has been approved as a monotherapy for adults with relapsed or refractory AML with FLT3 mutation (Pulte et al. 2021). Although FLT3 inhibitors have significantly improved the survival of FLT3-mutated AML patients, primary therapeutic resistance and short-lived responses remain still as a frequent problem.

Resistance to FLT3 inhibitors can arise from specific FLT3-TKD mutations or mutations in other oncogenes (McMahon et al. 2019, Alotaibi et al. 2021, Rummelt et al. 2021, Smith et al. 2022), resulting in activation of proliferation and pro-survival signaling pathways. Nonetheless, emerging evidence indicates that non-genetic factors, such as epigenetic, transcriptional, and metabolic states, also play a crucial role in resistance to chemotherapy and targeted therapies in AML (Bell et al. 2019, Petti et al. 2022). Recent studies propose that early resistance to cancer therapies might also be mediated by non-genetic mechanisms, particularly through drug-tolerant persister (DTP) states encoded by pre-existing intrinsic transcriptional heterogeneity (Shaffer et al. 2017, Cotton et al. 2023, Pellecchia et al. 2024). These DTP cells, closely resembling clinical measurable residual disease (MRD), can subsequently undergo further evolution to acquire diverse drug-resistance mechanisms, including mutations (Ramirez et al. 2016). Novel therapeutic strategies aimed at targeting drug-tolerant persister cells are crucial to prevent the development of drug resistance and achieve MRD-negative complete remission, thereby reducing relapse rates and prolonging survival. However, the mechanisms behind this early resistance to FLT3 inhibitors in AML patients remain poorly understood.

In this study, we employed our recently established lineage-tracing method ReSisTrace (Dai et al. 2024) to investigate the transcriptional states preceding resistance against FLT3 inhibitors midostaurin and quizartinib in an FLT3-ITD-positive MOLM-13 cell line. We show that targeting a pre-resistance signature gene G1 to S phase transition 1 (*GSPT1*) with a selective degrader (CC-90009) exhibits strong synergy when combined with FLT3 inhibitors in FLT3-ITD-mutated AML cell lines and primary patient samples. We further searched the L1000 database (Subramanian et al. 2017) for compounds inducing gene expression changes opposite to the pre-resistance signatures. Linsitinib (IGF1R and Src family kinase inhibitor), vistusertib (mTOR inhibitor), and meisoindigo (Src family kinase inhibitor), all targeting pathways downstream of or parallel to oncogenic FLT3 signaling, were predicted and validated to pre-sensitize FLT3-ITD-positive cell lines and primary patient samples to FLT3 inhibitors. In conclusion, our study demonstrates the effectiveness of ReSisTrace in uncovering targetable, pre-existing transcriptional features of treatment resistance in hematological cancers. Importantly, the approach allowed us to identify synergistic and pre-sensitizing drugs that have the potential to increase FLT3 inhibitor treatment efficacy in FLT3-ITD-positive AML by preventing emergence of treatment resistance.

## Results

### Establishing primed resistance signatures using lineage tracing

To elucidate transcriptional cell states that are primed to resist treatment with the FLT3 inhibitors quizartinib and midostaurin, we applied ReSisTrace in an FLT3-ITD-mutated AML cell line MOLM-13. In this approach, uniquely labeled and synchronized cells undergo one duplication before being randomly divided into two populations. One population is subjected to single-cell RNA-sequencing (scRNA-seq) analysis directly, while the other undergoes drug treatment (Fig. 1A). Cells subjected to drug treatment were exposed to either 100 nM midostaurin or 10 nM quizartinib for 72 hours to kill ∼70% of the cells. Subsequently, the surviving cells were allowed to recover, and scRNAseq-analysis was performed to identify the drug-resistant lineages within the post-treatment samples. The lineage information was used to annotate corresponding sister cells in the pre-treatment sample as pre-resistant cells. Cell lineages present in the pre-treatment sample but absent in the post-treatment sample were annotated as pre-sensitive. We further analyzed a control sample treated with DMSO only, mimicking the growth conditions of the drug treatment experiments, to differentiate between transcriptional states associated with experiment-specific growth fitness and those specifically linked to primed resistance.

**Fig. 1.**
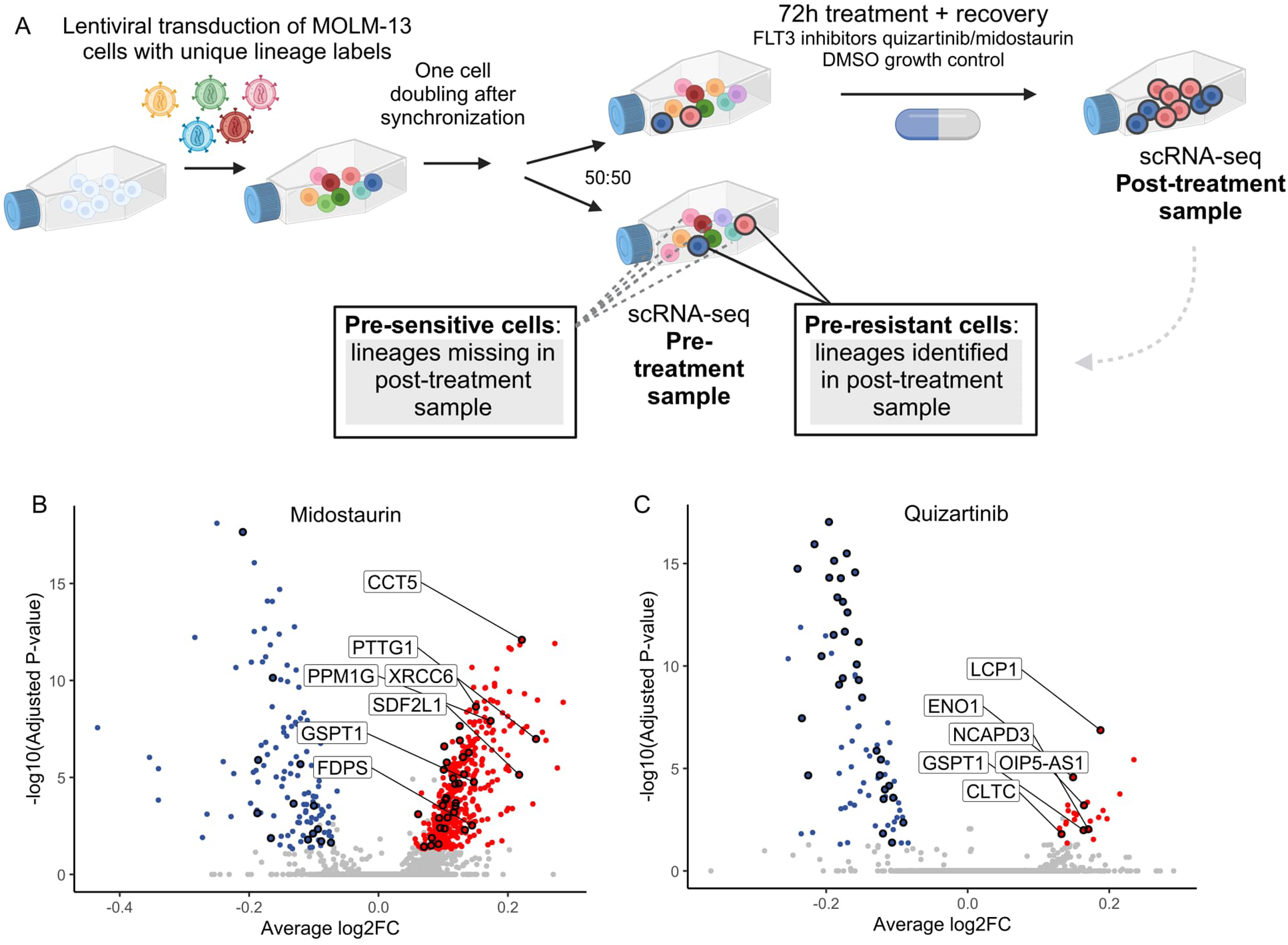
Utilization of ReSisTrace approach to study transcriptional states preceding resistance against FLT3 inhibitors in acute myeloid leukemia. A) Schematic representation of applying ReSisTrace approach to study primed resistance against FLT3 inhibitors in acute myeloid leukemia. Uniquely labeled, synchronized cells were permitted to duplicate once and were then divided into two groups for scRNA-sequencing (pre-treatment sample) and drug treatment. The lineage labels of cells that survived the treatment were determined by scRNA-sequencing (post-treatment sample) and were used to identify pre-resistant cells in the pre-treatment sample. (B-C) Midostaurin (B) and quizartinib (C) pre-resistance signatures were determined by comparing the gene expression profiles between the pre-resistant and pre-sensitive cells in the pre-treatment sample. Sister-concordant differentially expressed genes are shown in color. Genes differentially expressed specifically in the pre-resistance signatures compared to the pre-fitness signature obtained from the DMSO-treated growth control experiment are circled. Genes with the highest log2FC are labeled.

In pre- and post-treatment scRNA-seq samples, lineage labels consisting of 20 random bases were detected in approximately 85% of the cells passing quality control (Fig. S1A). On average, 71% of the lineages were unique for each treatment (Fig. S1B). In the pre-treatment samples, approximately 88% of the cells with lineage labels represented lineages with only one cell (Fig. S1C). We used only these cells in the downstream analyses to exclude lineages with increased basal growth rate and to reduce the rate for false-negative findings due to both sister cells ending up in the pre-treatment sample because of random sampling. We further validated that cells sharing the same lineage labels, i.e. sister cells, have significantly more similar transcriptomes than random cell pairs (Fig. S1D), suggesting that the sister cells can be used as proxies to study lineage-specific treatment resistance, in line with our results in the solid cancer setting (Dai et al. 2024).

We then explored the pre-treatment samples in the transcriptomic space. In the UMAP embedding, all cells within the pre-treatment samples were clustered together, indicating that the samples destined to undergo different treatments showed similar composition (Figs. S1E and S1F). Notably, no distinct subclones were found, indicating the homogeneity of the MOLM-13 cell line. Next, in the midostaurin pre-treatment sample, we determined 544 lineages as pre-resistant and 2015 lineages as pre-sensitive, based on the lineage labels observed or lacking in the post-treatment sample, respectively. Similarly, in the quizartinib pre-treatment sample, we annotated 344 lineages as pre-resistant and 3848 lineages as pre-sensitive. In the control pre-treatment sample, we found 801 lineages that remained in the respective post-treatment sample, and thus annotated as pre-fit cells, while 2840 lineages were annotated as pre-unfit cells. A UMAP projection did not reveal a clear separation between the pre-resistant and pre-sensitive cells, nor the pre-fit and pre-unfit cells (Figs. S1F-I), suggesting that there were no discernible subclonal patterns indicative of resistance.

We then performed differential gene expression analysis between the pre-resistant and pre-sensitive cells using the non-parametric Wilcoxon rank sum test, to obtain the midostaurin and quizartinib pre-resistance signatures (Figs. 1B and 1C; full gene lists are shown in Tables S1 and S2). Additionally, we compared the expression profiles of the pre-fit and pre-unfit cells in the DMSO-treated growth control experiment to elucidate the pre-fitness signature (Table S3). To determine the potential pathways associated with primed FLT3 inhibitor resistance, we ran gene set enrichment analysis (GSEA) on the pre-resistance and pre-fitness signature gene lists, ranked by adjusted *P*-value, *P*-value, and log2 fold change (log2FC). We found that many KEGG pathways linked to cell cycle, DNA replication, and DNA repair were enriched among the genes up-regulated in both pre-resistant and pre-fit cells (full lists of significant KEGG pathways are shown in Table S4). Moreover, KEGG pathways related to carbohydrate metabolism, including fructose and mannose metabolism, inositol phosphate metabolism, pentose phosphate pathway, as well as amino sugar and nucleotide sugar metabolism, were enriched among genes specifically up-regulated in the pre-resistant cells (Fig. 2A). When comparing the midostaurin and quizartinib pre-resistance signatures, we found that the Wnt signaling pathway was up-regulated in both of them, consistent with previous findings that activation of Wnt/beta-catenin signaling decreases the potency of FLT3 inhibitors in FLT3-ITD-mutated cells (Jiang and Griffin 2010). In contrast, phosphatidylinositol signaling system and mTOR signaling pathway were up-regulated only in the quizartinib pre-resistance signature. The analysis of the BioCarta pathways also indicated the up-regulation of mTOR signaling in quizartinib pre-resistance signature, while MAPK and ERK signaling pathways were up-regulated in the midostaurin pre-resistance signature (Fig. 2B; full lists of significant BioCarta pathways are shown in Table S5).

**Fig. 2.**
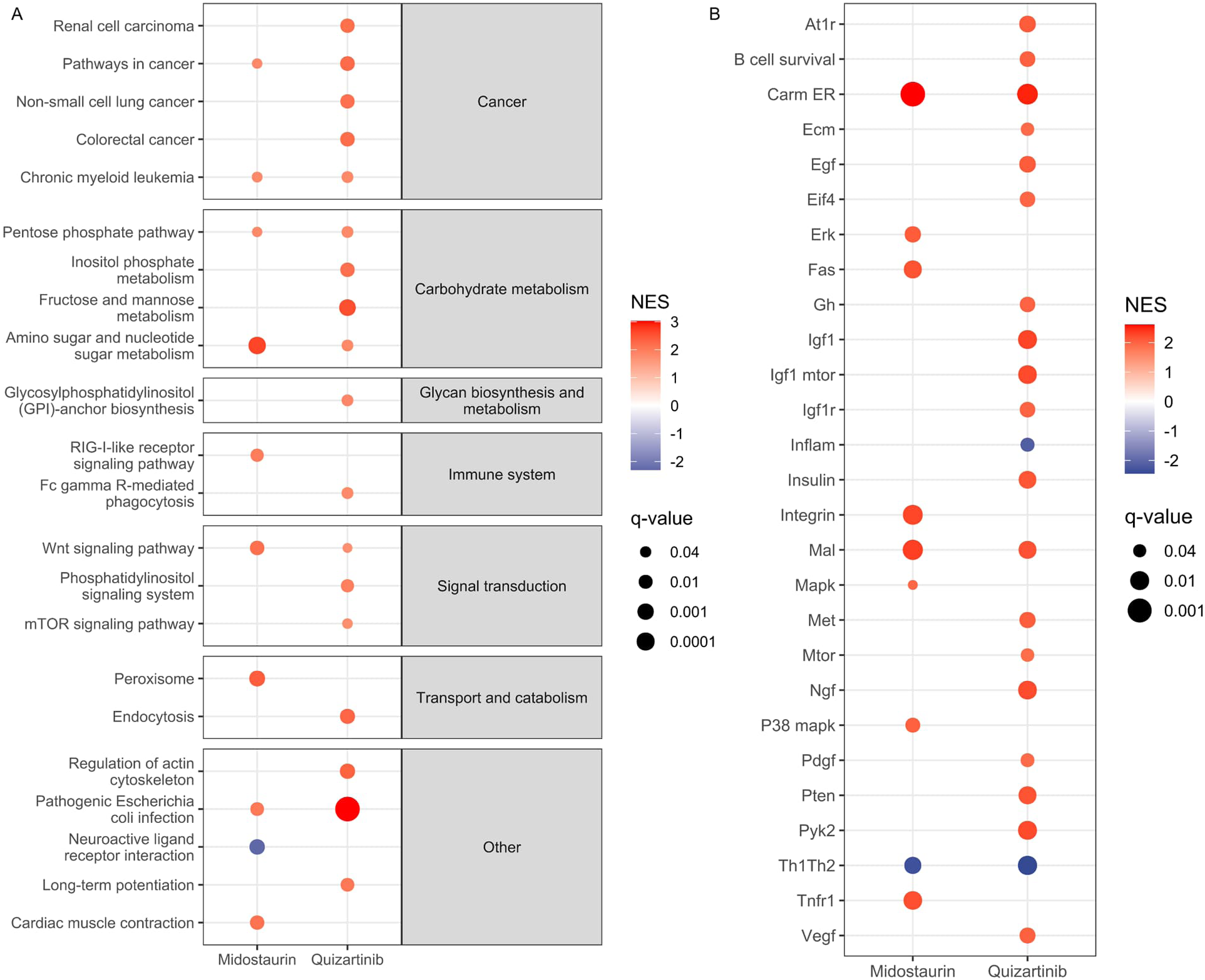
Gene set enrichment analysis. (A) KEGG and (B) BioCarta pathways specifically enriched among the midostaurin or quizartinib pre-resistance signature genes.

### Targeting individual pre-resistance signature genes with inhibitors

As we observed an overlap between the top up-regulated genes in the pre-resistance signatures and those in the growth control (pre-fit versus pre-unfit cells), we sought to identify genes exhibiting increased expression specifically within the pre-resistance signatures compared to the pre-fitness signature. To accomplish this, we employed permutation and bootstrapping tests to assess the significance of the fold change differences between the signatures. Our analysis revealed that 41 genes were significantly (adjusted *P*-value < 0.05 in pre-resistant vs pre-sensitive) and specifically (permutation and bootstrapping test between pre-resistance and pre-fitness log2FC, *P* < 0.05) up-regulated in midostaurin pre-resistant cells, and 7 genes in quizartinib pre-resistant cells when compared to both their respective pre-sensitive cells and the pre-fit signal within the DMSO control (Figs. 1B and 1C; Tables S6 and S7; full results are shown in Tables S1 and S2).

We selected one gene, G1 to S phase transition 1 (*GSPT1*), which was the only targetable gene present among the top genes in the pre-resistance signatures for both midostaurin and quizartinib treatments (Tables S6 and S7), to investigate whether combining a GSPT1 inhibitor with FLT3 inhibitors could enhance treatment efficacy. *GSPT1* encodes a protein involved in translation termination, and it could be targeted for selective ubiquitin-mediated degradation with a molecular glue CC-90009 (Hansen et al. 2021). Our experimental approach involved pre-treating MOLM-13 cells with CC-90009 for 24 hours, followed by the addition of midostaurin or quizartinib for an additional 72 hours. CC-90009 showed synergy with both midostaurin and quizartinib according to the common reference models, including Bliss (Bliss independence), HSA (Highest Single Agent), Loewe (Loewe additivity), and ZIP (Zero Interaction Potency) (Figs. 3A, 3B, and 3D; dose-response matrices in Figs. S2A and S2B). Importantly, the combination was synergistic also in another FLT3-ITD-positive cell line, MV4-11 (Figs. 3E, 3F, 3H, S2D, and S2E). To confirm the expected mechanism, we verified by western blotting that CC-90009 treatment led to the degradation of GSPT1 in both MOLM-13 and MV4-11 cells (Fig. S3). In MOLM-13, we observed predominantly low expression of the long isoform (∼80 kDa) and elevated levels of a shorter (∼40 kDa) isoform/fragment. Following CC-90009 treatment, there was a 73% to 80% reduction in the shorter isoform/fragment compared to the DMSO-treated control (Fig. S3A). In turn, MV4-11 cells exhibited elevated expression of the long GSPT1 isoform, accompanied by the presence of the shorter fragment. Both isoforms were significantly reduced by 39% to 58% after CC-90009 treatment (Fig. S3B). Taken together, our results show that CC-90009 synergizes with the FLT3 inhibitors midostaurin and quizartinib by decreasing GSPT1, and that these combinations could potentially be used to increase treatment efficacy in FLT3-ITD-mutated AML.

**Fig. 3.**
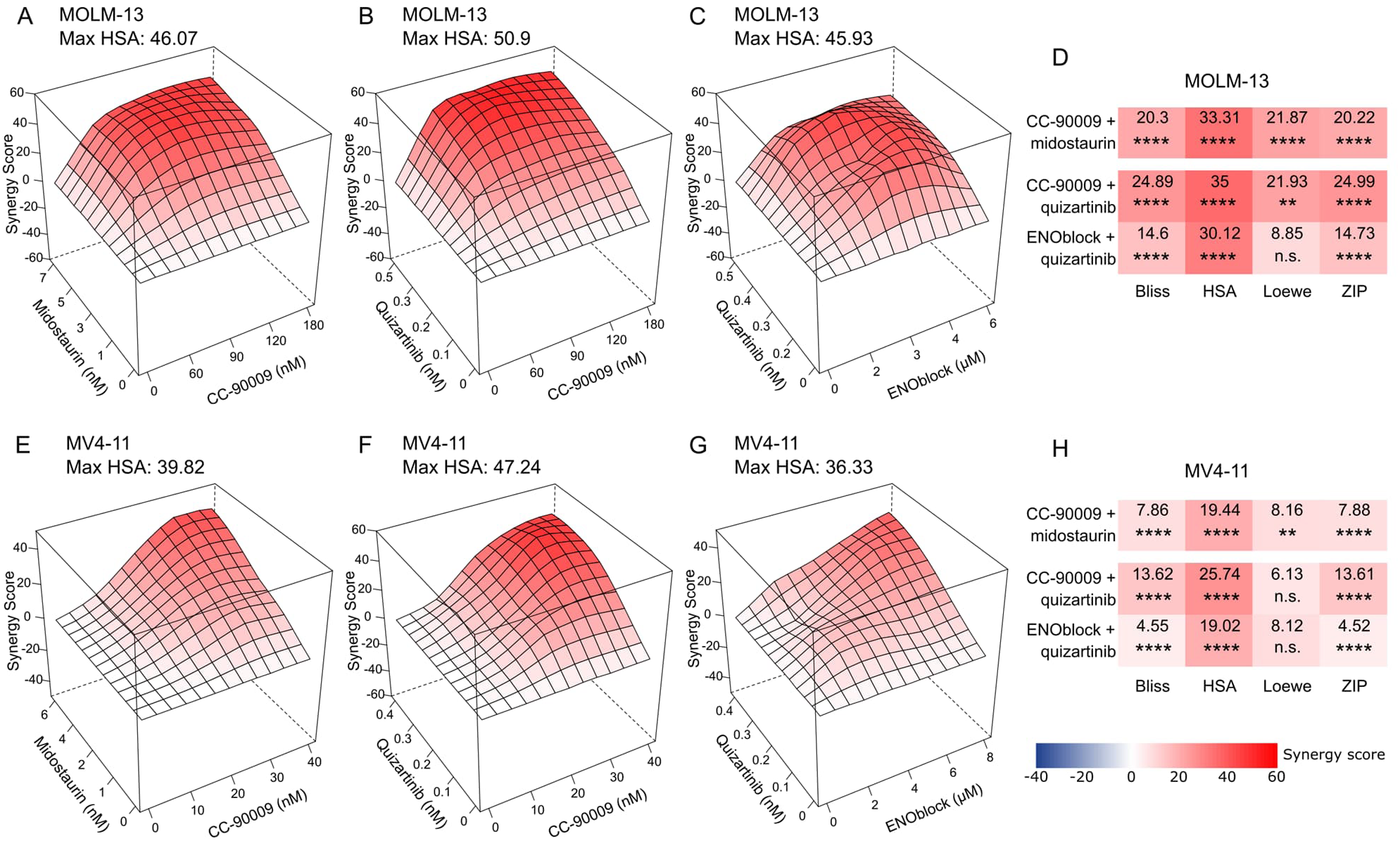
Synergistic effect of drugs targeting individual pre-resistance genes in combination with FLT3 inhibitors midostaurin or quizartinib. HSA synergy landscapes (A-C, E-G) and mean synergy scores (D and H) for CC-90009 and ENOblock in combination with midostaurin or quizartinib in MOLM-13 (A-D) and MV4-11 cells (E-H). (D and H) *P*-values were determined by bootstrapping test; n.s. = not significant, \**P* < 0.05, \*\**P* < 0.01, \*\*\**P* < 0.001, \*\*\*\**P* < 0.0001.

In our search for specific combination therapies for quizartinib, we further selected a top gene enolase 1 (*ENO1*) from the pre-resistance signature (Table S7). *ENO1* is a gene encoding a glycolytic enzyme alpha-enolase, which can be targeted by ENOblock (Jung et al. 2013). Similarly, we pre-treated the cells with ENOblock before adding quizartinib. In both MOLM-13 and MV4-11 cell lines, ENOblock showed synergy with quizartinib according to Bliss, HSA, and ZIP models (Figs. 3C, 3D, 3G, and 3H; dose-response matrices in Figs. S2C and S2F). To study whether the synergistic effect of ENOblock with quizartinib could result from the inhibition of the enzymatic activity or only the non-glycolytic functions of enolase (Jung et al. 2013, Satani et al. 2016, Cho et al. 2017), we investigated the synergistic potential of sodium fluoride, a known inhibitor of enolase enzymatic activity, when combined with quizartinib treatment. In MOLM-13, sodium fluoride failed to show significant synergism with quizartinib treatment (Figs. S4A, S4C, and S4D). In MV4-11, while the HSA score indicated moderate synergism (Figs. S4B, S4C, and S4E), other models demonstrated negative synergy scores (Fig. S4C). These results suggest that the non-glycolytic functions of enolase, inhibited by ENOblock, may play a more pivotal role in priming resistance against quizartinib compared to the enolase enzymatic activity.

### Targeting pre-resistant states by drugs with opposite gene expression signatures

In addition to targeting individual genes in the pre-resistance signature, we employed a systematic strategy to target primed resistance in a more comprehensive manner. This involved the use of small molecules that would shift cancer cells toward the pre-sensitive states, thereby enhancing their responsiveness to FLT3 inhibitor treatment. To this end, we queried the L1000 database (Subramanian et al. 2017) for drugs that would induce gene expression changes opposite to the pre-resistance signatures (i.e. similar to the pre-sensitivity signatures). To obtain more robust results, we used multiple variants of the pre-resistance signatures, two of which were adjusted for gene expression patterns associated with experimental growth fitness (see Methods). The drugs were filtered by negative connectivity scores with *P*-value < 0.05 across all the pre-resistance signature variants (Figs. 4A and 4B; Table S8). Notably, some of the identified drugs target kinases downstream of FLT3, such as vistusertib, which inhibits mTOR, GDC-0068 inhibiting AKT, and meisoindigo inhibiting multiple SRC family kinases along with IGF1R. Additionally, some of the identified drugs inhibit kinases activating the same signaling pathways as FLT3, such as linsitinib targeting IGF1R and insulin receptor, and CGP-52411 targeting EGFR. Among the other potential pre-sensitizing drugs were CDK inhibitors, a CHEK1 inhibitor, and OTS167, which targets MELK and has been reported to inhibit FLT3 protein translation and synergize with the FLT3 inhibitor gilterinib in FLT3-ITD-mutated AML (Eisfelder et al. 2021). When combined with midostaurin, linsitinib (IGFR1/insulin receptor inhibitor) and vistusertib (mTOR inhibitor) demonstrated synergism across all four synergy models (Figs. 4C, 4D, and 4H; Dose-response matrices in Figs. S2G, and S2H). In turn, in combination with quizartinib, AZD5438 (CDK inhibitor) displayed synergism only according to the HSA model (Figs. 4E, 4H, and S2I), while OTS167 (MELK inhibitor) exhibited additive effect or low synergism according to three models (Figs. 4G, 4H, and S2K). Notably, meisoindigo (SRC family kinase and IGF1R inhibitor) demonstrated higher synergism with quizartinib according to Bliss, HSA, and ZIP models (Figs. 4F, 4H, and S2J). We then evaluated the effect of the combinations, which showed the highest synergism, in the other FLT3-ITD-positive cell line, MV4-11. We found that vistusertib consistently displayed the highest synergism with midostaurin (Figs. 4J, 4L, and S2M), while linsitinib with midostaurin and meisoindigo with quizartinib demonstrated synergism according to HSA and Loewe models (Figs. 4I, 4K, 4L, S2L, and S2N). Interestingly, all the top-performing predicted drugs, linsitinib, vistusertib and meisoindigo, inhibit pathways downstream or parallel to FLT3 signaling (Fig. 4M). Our results indicate that these pathways are activated in the pre-resistant cells and inhibiting these pathways pre-sensitize FLT3-ITD-mutated cells to FLT3 inhibitors.

**Fig. 4.**
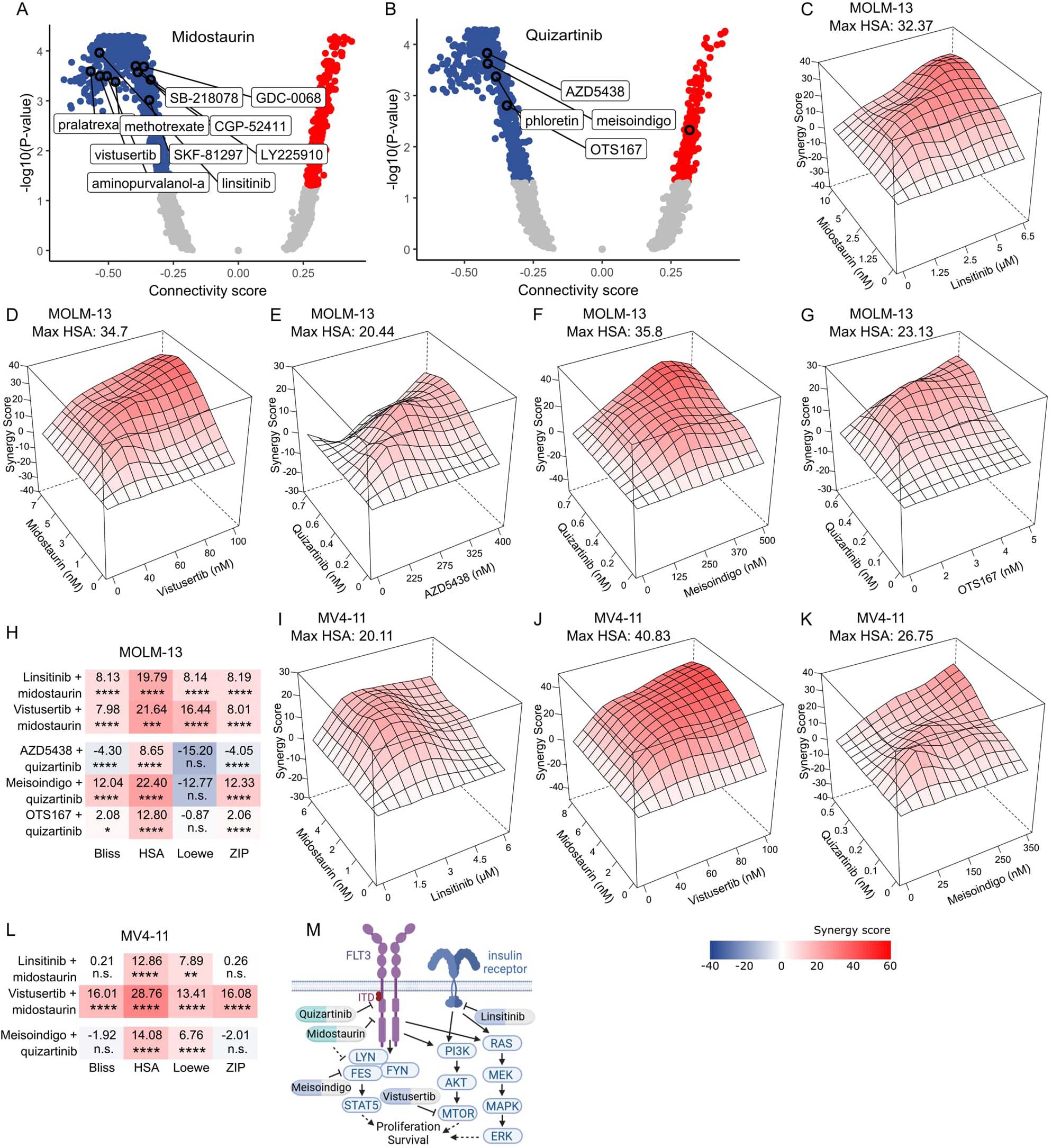
Targeting pre-resistant states with drugs with opposite gene expression signatures. (A-B) Connectivity scores between the L1000 drug perturbed signatures and unfiltered (A) midostaurin and (B) quizartinib pre-resistance signatures. Drugs with significant connectivity scores in all signature variants are circled. (C-L) HSA synergy landscapes (C-G, I-K) and mean synergy scores (H and L) for the indicated drugs in combination with midostaurin or quizartinib in MOLM-13 (C-H) and MV4-11 (I-L) cells. (H and L) *P*-values were determined by bootstrapping test; n.s. = not significant, \**P* < 0.05, \*\**P* < 0.01, \*\*\**P* < 0.001, \*\*\*\**P* < 0.0001. (M) Schematic representation of pathways targeted by the top-performing pre-sensitizing drugs.

### Validating drug combinations in FLT3-ITD-mutated patient samples

We then evaluated the efficacy of the drug combinations in three diagnosis-stage FLT3-ITD-mutated AML patient samples (Fig. 5A), each with blast cell percentage ≥ 58% and an FLT3-ITD variant allele frequency ≥ 69% (Table S9). Initially, we cultured the samples in both Mononuclear Cell Medium (MCM) and 25% HS-5 stromal cell-conditioned RPMI-1640 medium. Despite previous reports showing that HS-5 cell-conditioned medium protects FLT3-ITD-mutated primary AML cells from FLT3 inhibitors (Karjalainen et al. 2017), we chose this medium for our experiments as it more closely mimics the in vivo environment and supported the growth of the primary cells. MCM, in contrast, did not support primary cell growth for the required 4 days (Fig. S5A).

**Fig. 5.**
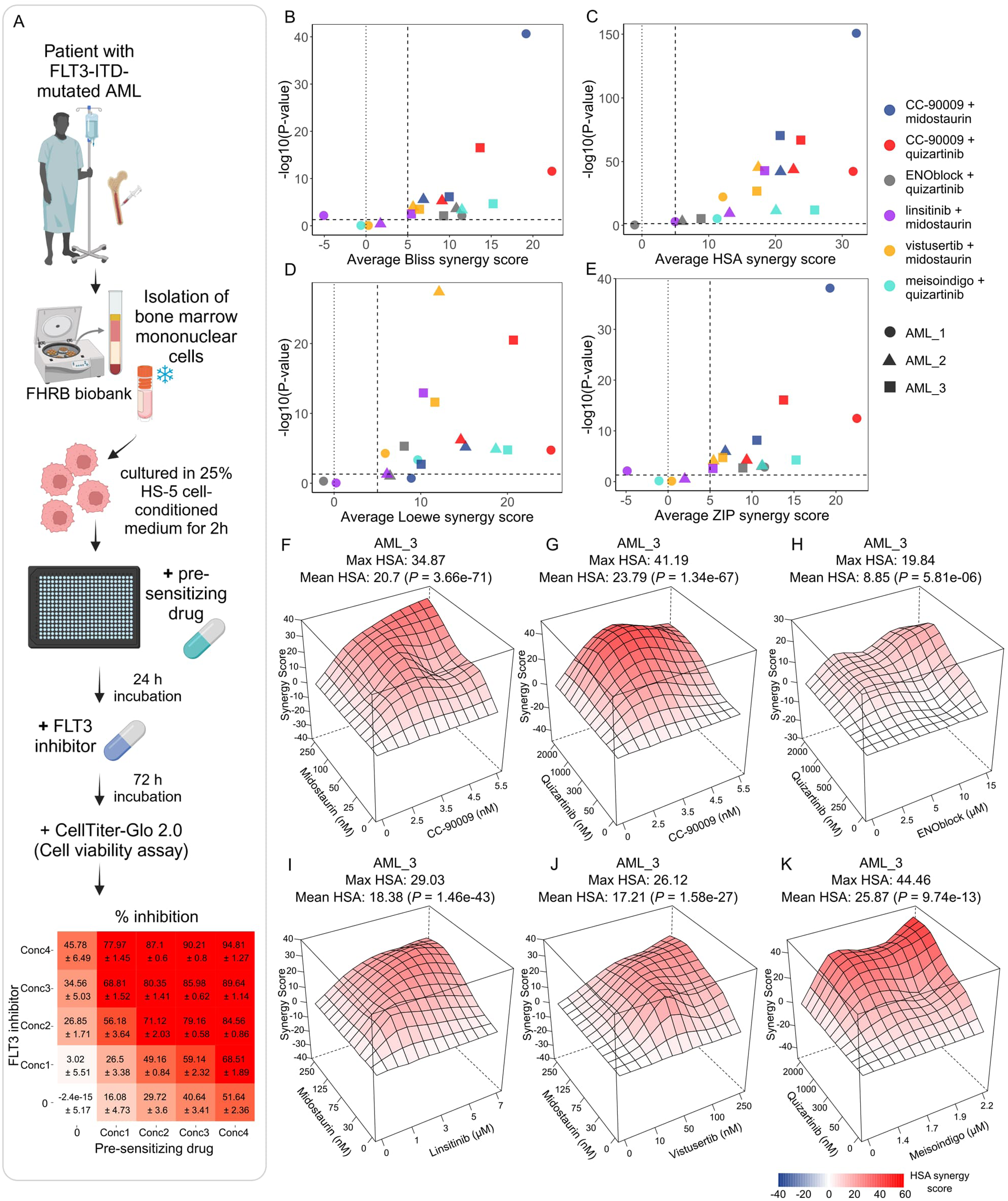
Validation of drug combinations in patient samples. (A) Schematic representation of testing drug combinations in primary AML patient samples. (B-E) Average synergy scores according to different synergy models for the six drug combinations in three FLT3-ITD-mutated AML patient samples. (F-K) HSA synergy landscapes for the six drug combinations in patient sample AML_3. *P*-values were determined by bootstrapping test.

In single-drug treatments, midostaurin and meisoindigo were less effective in primary AML samples compared to MOLM-13 and MV4-1 cell lines. Interestingly, primary cells were more sensitive to CC-90009 than the cell lines (Figs. S2 and S5). Quizartinib did not achieve 50% inhibition in the primary AML cells even with the highest concentration tested (2 µM), while ENOblock was either ineffective or promoted growth at concentrations up to 15 µM (Fig. S5). Among the drug combinations, CC-90009 consistently exhibited high synergy with quizartinib across all three samples, as confirmed by all four synergy models (Fig. 5B-5E, 5G, S6B, and S6H; dose-response matrices in Figs. S5C, S5I, and S5O). Additionally, CC-90009 combined with midostaurin showed strong synergy across all samples, with only the Loewe synergy score in sample AML_1 failing to reach statistical significance (Figs. 5B-5F, S6A, and S6G; dose response Figs. S5B, S5H, and S5N). Vistusertib combined with midostaurin, and meisoindigo with quizartinib showed synergism in all samples according to HSA and Loewe models, and in two samples (AML_2 and AML_3) according to Bliss and ZIP models (Figs. 5B-5E, 5J, 5K, S6E, S6F, S6K, and S6L; dose response Figs. S5F, S5G, S5L, S5M, S5R, and S5S). Linsitinib and midostaurin combination displayed significant synergy scores according to all models but only in sample AML_3 (Figs. 5B-5E, and 5I; dose response Fig. S5Q) suggesting that certain resistance-conferring pathways may be activated in a subset of patients. Meanwhile, ENOblock combined with quizartinib showed synergy in AML_2 and AML_3 (Figs. 5B-5E, 5H, and S6I) but the overall inhibition levels remained low, likely due to the ineffectiveness of both drugs as single agents (Fig. S5J and S5P).

Taken together, targeting GSPT1 with CC-90009 exhibits strong synergy with FLT3 inhibitors in both FLT3-ITD-positive cell lines and primary AML patient samples. This combination represents a promising approach to enhance treatment efficacy in FLT3-ITD-mutated AML. Furthermore, the three top-performing drugs, vistusertib, linsitinib and meisoindigo, predicted to pre-sensitize cells to FLT3 inhibitors, demonstrated synergy with the FLT3 inhibitors in both FLT3-ITD mutated patient samples and cell lines. These findings validate the effectiveness of our ReSisTrace lineage-tracing method for identifying new, rational drug combinations targeting primed treatment resistance in hematological cancers.

## Discussion

Despite considerable progress in developing more potent and selective FLT3 inhibitors, the emergence of resistance persists as a significant challenge. Studies investigating AML patient relapses after FLT3 inhibitor therapy have consistently identified alterations in genes within the RAS/MAPK pathway, particularly NRAS and KRAS, as the most prevalent genetic aberrations associated with acquired resistance, alongside novel FLT3 mutations (McMahon et al. 2019, Alotaibi et al. 2021, Smith et al. 2022). Moreover, FLT3-independent activation of RAS/MAPK and/or PI3K/AKT/mTOR signaling pathways, along with the sustained expression of genes involved in FLT3-mediated cellular transformation, has been observed in FLT3 inhibitor-resistant cell lines and primary samples (Piloto et al. 2007, Lindblad et al. 2016). Additionally, non-genetic mechanisms, such as soluble factors from the bone marrow microenvironment, can activate the downstream RAS/MAPK, PI3K/AKT, and mTOR signaling in AML cells, contributing to early treatment resistance against FLT3 inhibitors (Traer et al. 2016, Joshi et al. 2021, Park et al. 2022). Furthermore, intrinsic transcriptional heterogeneity can also induce drug tolerance in cancer cells (Shaffer et al. 2017). Our ReSisTrace lineage-tracing method offers a means to unbiasedly detect pre-existing transcriptional states leading to resistant fates. Applying the method on FLT3-ITD-positive cells unveiled transcriptional states characterized by increased signaling of the aforementioned resistance-conferring pathways, priming cells for resistance against FLT3 inhibitors within the treatment-naïve cell population. Reverting these cell states that are predisposed to survive the treatment, already before more stable resistance emerges, could represent a viable strategy to enhance long-term remission rates in FLT3-ITD-positive AML patients.

Our lineage-tracing method not only allows characterization of resistant populations prior to the treatment, but also reveals the vulnerable states that could be induced via pre-treatments. We targeted primed resistance with drugs that were predicted to pre-sensitize cells to FLT3 inhibitors based on the similarity between our pre-sensitivity signature and the L1000 drug-perturbed consensus signatures. Among these drugs, many target pathways that are parallel to or downstream of oncogenic FLT3 signaling (Fig. 4O). For instance, vistusertib inhibits mTOR activity, attenuating any FLT3-ITD-independent resistance-promoting mTOR signaling when combined with an FLT3 inhibitor. Consistent with this, a dual PI3K/mTOR inhibitor PF-04691502 has been found to display synergistic cytotoxicity with quizartinib in FLT3-ITD AML cells (Darici et al. 2021). Another predicted drug, linsitinib, selectively targets IGF1R and insulin receptor (INSR) (Mulvihill et al. 2009). Notably, only INSR is moderately expressed in the MOLM-13 and MV4-11 cell lines, according to our scRNA-seq data and the DepMap Portal (https://depmap.org/portal). Upon binding of insulin or IGF1, INSR activates multiple downstream pathways, including PI3K/AKT/mTOR and RAS/MAPK/ERK signaling (Hopkins et al. 2020). Similarly, meisoindigo inhibits IGF1R but not INSR (Tegethoff et al. 2017). Since IGF1R is not expressed in the MOLM-13 and MV4-11 cells, meisoindigo’s main mechanism of action in these cell lines likely involves inhibition of several non-receptor tyrosine kinases, including FES, FER, and the SRC family kinases LYN, FYN, YES1, LCK, and SRC (Tegethoff et al. 2017). Among these, LYN and FES are expressed in MOLM-13 and MV4-11 and play key roles in oncogenic FLT3 downstream signaling (Okamoto et al. 2007, Voisset et al. 2010). In primary AML cells, meisoindigo demonstrated strong synergy with quizartinib in two out of three samples, with synergy scores even higher than in the cell lines. Interestingly, meisoindigo has been used to treat chronic myeloid leukemia in China for decades (Xiao et al. 2002) and has also been reported to induce apoptosis in AML cell lines and primary AML cells (Lee et al. 2010). Notably, vistusertib and linsitinib, when combined with midostaurin, also showed enhanced synergy in the same two primary AML samples as meisoindigo. Together, these findings suggest that the resistance-conferring pathways targeted by these pre-sensitizing drugs may be active and more critical in a subset of FLT3-ITD-mutated AML patients. Further studies are needed to clarify the mechanisms driving the activation of these pathways and to better understand the precise mechanism of action of these drug combinations.

Among the drugs predicted to pre-sensitize cells to FLT3 inhibitors, some showed low synergy or even antagonistic effects when tested in MOLM-13 in combination with FLT3 inhibitors. This discrepancy might stem from decreased accuracy of predictions when comparing our pre-resistance signatures against the L1000 phase 1 consensus signatures, generated from drug-perturbed gene expression profiles of cell lines derived from solid cancers, not including data from blood cancer cell lines. Furthermore, the detection of subtle transcriptional changes that mediate primed resistance against FLT3 inhibitors, while considering the accompanying growth fitness-related alterations, presents several challenges. The up-regulation of pathways downstream of oncogenic FLT3 signaling could 1) confer resistance to the FLT3 inhibitors, indicated by our GSEA findings and pathways targeted by the validated pre-sensitizing drugs, and 2) promote increased proliferation and fitness in the non-treatment control, given that FLT3-ITD acts as the oncogenic driver of MOLM-13 cells. This dual effect complicates the distinction between resistance-related alterations and those providing growth advantage. Additionally, cell cycle synchronization via thymidine block introduces stress and imposes a non-random selection pressure to all samples, potentially inducing noise that obscures the pre-resistance signal, and thus warrants careful consideration when analyzing sensitive cancer models.

As a model for FLT3-ITD-mutated AML, MOLM-13 cell line is heterozygous for the FLT3-ITD mutation, also expressing the wild-type FLT3 receptor. It has been shown that the ligand-dependent activation of wild-type FLT3 is only minimally affected by quizartinib and midostaurin (Kawase et al. 2019), and that the co-existence of wild-type FLT3 can dampen the efficacy of FLT3 inhibitors in FLT3-mutated AML cells in vitro and in vivo (Chen et al. 2016). As a result, signaling through wild-type FLT3 in MOLM-13 cells could contribute to resistance mechanisms not observed in FLT3-ITD^+/+^ AML cells. However, most drug combinations that demonstrated synergistic effects in MOLM-13 cells also showed synergy in the homozygous FLT3-ITD^+/+^ MV4-11 cell line, as well as in FLT3-ITD-mutated primary AML patient samples with a high FLT3-ITD variant allele frequency, indicative of a homozygous mutation. These findings suggest that the drug combinations are effective in both genetic backgrounds and that the underlying primed resistance mechanisms are not unique to the MOLM-13 cell line.

We observed an up-regulation of GSPT1 in both midostaurin and quizartinib pre-resistance signatures. CC-90009, a selective degrader of GSPT1, has emerged as a promising candidate for novel AML treatment, showing efficacy across various genetic backgrounds (Hansen et al. 2021). Ongoing clinical trials are assessing CC-90009’s potential, including a phase 1 dose-finding study in relapsed or refractory patients (ClinicalTrials.gov: NCT02848001), and a phase 1b study evaluating the safety and efficacy of CC-90009 in combination with gilterinib (an FLT3 inhibitor), venetoclax (BCL2 inhibitor), or the nucleoside analog azacytidine (ClinicalTrials.gov: NCT04336982). Depletion of GSPT1 through drug-induced degradation leads to impaired translation termination, activation of the integrated stress response pathway, and TP53-independent cell death in AML cells, with minimal effect on normal hematopoietic stem cells (Sellar et al. 2022). Further, GSPT1 depletion attenuates translation initiation and cell cycle progression by inhibiting mTOR signaling downstream of AKT activation, independent of the Erk1/2 pathway (Chauvin et al. 2007). Whether increased GSPT1 expression directly up-regulates mTOR signaling and partly explains how GSPT1 could confer resistance against FLT3 inhibitors, warrants further investigation. Interestingly, hyperactivation of the mTOR signaling pathway has been found to attenuate the response to CC-90009 by blocking GSPT1 degradation (Surka et al. 2021). Therefore, the synergistic effect of CC-90009 and quizartinib or midostaurin may partly stem from FLT3 inhibitors reducing downstream mTOR signaling, thus enhancing the efficacy of CC-90009 in FLT3-ITD-positive AML cells. Notably, while quizartinib alone was largely ineffective in primary AML patient samples when tested in clinically relevant concentrations (peak serum concentration range from 250 nM during induction therapy to 945 nM at maintenance therapy with VANFLYTA®; https://www.accessdata.fda.gov/drugsatfda_docs/label/2023/216993s000lbl.pdf), it, like midostaurin, exhibited strong inhibitory effects across all samples when combined with low concentrations of CC-90009. This highlights the potential of combining CC-90009 with FLT3 inhibitors to enhance treatment efficacy while reducing the toxicities and side-effects associated with high-dose FLT3 inhibitor treatments.

In summary, ReSisTrace lineage-tracing offers a powerful tool for elucidating intricate cellular states that prime treatment resistance also in hematological cancers, as demonstrated here in the context of FLT3-mutated AML. Furthermore, our study unveils novel synergistic and pre-sensitizing drugs that could potentially be used to increase FLT3 inhibitor treatment efficacy and prevent emergence of treatment resistance against FLT3 inhibitors in FLT3-ITD-positive AML.

## Methods

### Cell culture

AML cell lines MOLM-13 (kind gift from prof. Caroline Heckman; authentication of the cell line was performed with the Promega GenePrint24 System) and MV4-11 (CRL-9591, ATCC), as well as the bone marrow mesenchymal stromal cell line HS-5 (kind gift from prof. Caroline Heckman) were grown in RPMI-1640 (31870025, Gibco) supplemented with 10% heat-inactivated fetal bovine serum (FBS) (10500064, Gibco), 1% penicillin-streptomycin (15140122, Gibco), and 1% GlutaMAX (35050038, Gibco). Conditioned medium used for culturing primary AML patient samples was collected from 60% to 80% confluent HS-5 cell cultures after 72 hours of incubation, cleared by centrifugation, filtered through a 0.22-µm filter, and stored at -80°C. Cell lines were tested regularly for mycoplasma contamination with Venor GeM Classic mycoplasma detection kit (Minerva Biolabs, Berlin, Germany).

### Patient sample processing

Fresh-frozen bone marrow mononuclear cells from three FLT3-ITD-mutated AML patients at diagnosis, pre-treatment phase were obtained from the Finnish Hematology Registry and Clinical Biobank (FHRB) with appropriate ethical approval. Patient characteristics are described in Table S9. Samples were selected based on high blast cell percentage and a high variant allele frequency of the FLT3-ITD mutation. Following thawing, cells were cultured for 2 to 3 hours in a mixture consisting of 25% HS-5 stromal cell-derived conditioned medium and 75% RPMI-1640, supplemented with 10% FBS, 1% penicillin-streptomycin, and 1% GlutaMAX, before seeding for drug combination testing. Mononuclear Cell Medium (MCM, C-28030, PromoCell, Heidelberg, Germany) was also evaluated as a culturing medium for the primary AML cells. Protocols for processing the samples were approved by the Ethics Committees of Helsinki University Central Hospital.

### Small molecules

CC-90009 (HY-130800), ENOblock (HY-15858), meisoindigo (HY-13680), pralatrexate (HY-10446), vistusertib (HY-15247), quizartinib (HY-13001) were purchased from MedChemExpress (Monmouth Junction, NJ, USA), and AZD5438 (S2621), methotrexate (S1210), midostaurin (S8064), and OTS167 (S7159) from Selleckchem (Houston, TX, USA). Linsitinib (CT-O906-2) was from ChemieTek (Indianapolis, IN, USA).

### ReSisTrace experimental workflow

Lineage barcode library (pBA439_UMI20) was synthesized, and lentiviruses were packaged as previously described (Dai et al. 2024). Two million MOLM-13 cells were transduced with the lentivirus library to obtain 20% transduction efficiency. Cells were centrifuged with the lentiviral particles at 800 x g for 30 minutes and incubated at normal growth conditions o/n, after which cells were washed and grown for 3 days before adding 0.5 ug/ml puromycin (sc-108071, Santa Cruz Biotechnology) for 9 days to remove cells without barcodes. Barcoded cells (1.2 x10^6^) were synchronized by incubating with 2 mM thymidine (T1895, Merck, Darmstadt, Germany) for 24 hours, after which the cells were released from the thymidine block by washing, and 20,000 cells/well in 50 ul of growth medium were seeded on round-bottom 96-well plates. After 24 hours of incubation, 4 of the replicate wells were used for cell counting to ensure that the cells have approximately doubled, and 3 of the replicate wells were used as samples for the subsequent experiment. Each of these 3 samples were resuspended, and half of each sample (∼20,000 cells) was analyzed by scRNA-seq as pre-treatment samples, while 25 ul of growth medium was added to the remaining half of each sample, after which the cells were treated with midostaurin (100 nM), quizartinib (10 nM), or DMSO (MP Biomedicals, 0.05%) for 72 hours. The drug concentrations were chosen to kill ∼70% of cells compared to DMSO-treated cultures. After the treatment, cells were washed 4 times, each time by adding 200 ul of medium, centrifuging the plates 300 x g 5 min, and removing 200 ul of medium. Cells were then allowed to recover for 4 (midostaurin), 3 (quizartinib), and 7 days (DMSO/growth control), after which ∼20,000 cells from each sample were analyzed by scRNA-seq as post-treatment samples.

### Single-cell RNA-sequencing library preparation

Single-cell RNA sequencing libraries were generated with 10x Genomics’ Chromium Next GEM Single Cell 3’ Kit v3.1 (PN-1000269, 10x Genomics, Pleasanton, CA, USA), Chromium Next GEM Chip G Single Cell Kit (PN-1000127), and Dual Index Kit TT Set A (PN-1000215). Libraries were quantified with Agilent TapeStation (High Sensitivity D5000 ScreenTape system, Agilent Technologies, Santa Clara, CA, USA) and qPCR (KAPA Library quantification kit KK4835, Roche, Basel, Switzerland), and sequenced with NovaSeq 6000 (Illumina, San Diego, CA, USA).

### Lineage label mapping and processing

The lineage barcode sequence (5’-CTGGGGCACAAGCTTAATTAAGAATTCANNNNTGNNNNACNNNNGANNNNGTNNNNCTAGGGCCTAGAGGGCCC GTTTAAAC-3’) was incorporated as an individual gene *pBA439_UMI_20* into the GDCh38.d1.vd1 human reference genome alongside GENCODE v25 annotation to build the reference for running the Cell Ranger (v 5.0.1) (Zheng et al. 2017) pipeline to perform alignment and UMI quantification using a customized reference.

From the “possorted_genome_bam.bam” file generated by the “cellranger count” command, we extracted lineage label sequences for each cell. Specifically, we considered sequences mapped to the *pBA439_UMI_20* gene and the sub-pattern “CANNNNTGNNNNACNNNNGANNNNGTNNNNCT.” To account for sequencing errors, we applied the directional network-based method from UMI-tools (v1.0.1) (Smith et al. 2017) to correct sequences that differed by a single base from the representative sequence. Additionally, we filtered out non-unique lineage label sequences expressed by more than four cells in the pre-treatment samples. We categorized cells from each pre-treatment sample into two groups: 1) pre-resistant cells with lineage labels matching those of corresponding post-treatment samples, and 2) pre-sensitive cells assigned to lineages not detected after treatment.

### Preprocessing of the scRNA-seq data

We used the Seurat (v4.0.4) (Hao et al. 2021) to perform the data quality control, normalization, top variable gene selection, scaling, dimensionality reduction, and differentially expressed gene (DEG) analysis. For each ReSisTrace sample, the genes expressed in less than three cells were removed. We filtered the cells based on the distribution of the UMI counts ([20000, 200000] for pre-treatment sample; [10000, 100000] for post-treatment sample), the number of genes ([2500, 12000] for pre-treatment sample; [2000, 10000] for post-treatment sample), and percentage of mitochondrial transcripts (less than 15%). We used the “NormalizeData” function (default parameter setting) to normalize gene counts.

For clustering purposes, we merged the pre-treatment samples, including lineages with only one cell. After normalization and finding the top 3,000 variable genes by using the “FindVariableFeatures” function (default parameter setting), we assigned each cell a cell-cycle score based on its expression of G2/M and S phase markers by using the “CellCycleScoring” function. Then, we used the “ScaleData” function to scale gene counts, regressing out multiple variables, including the cell cycle scores, number of UMIs per cell, and percentage of mitochondrial gene expression. We used 30 principal components for UMAP projection and clustering, with Louvain algorithm and resolution 0.5.

### Sister cell similarity and sister-concordant genes

We pooled the pre-treatment samples from all the experiments together and defined the cells with matching labels within each treatment group as sister cells. Lowly expressed genes whose total normalized expressions are less than 0.61 (10% quantile) were removed. The Euclidean distances from top 1000 variant genes were calculated to evaluate the similarity between 1905 sister cell pairs and 1.44 x 10^8^ random cell pairs. The similarities for all sister cell pairs and 100 000 randomly selected random cell pairs were visualized in Fig. S1D.

For extracting the sister concordant genes, we considered only the lineages with two sister cells, and removed the sister cells with the Euclidean distance of their transcriptomes larger than 17.86 (90% quantile). The genes with log2FCs significantly lower in the sister cell pairs compared to random cell pairs were defined as the sister-concordant genes (two-tailed t-test with an FDR-adjusted P value threshold of 0.05).

### Pre-resistance gene expression signatures

The “FindMarkers” function (test.use = “wilcox”, logfc.threshold = 0, min.pct = 0) from Seurat R package was used to determine the differentially expressed genes by Wilcoxon rank sum test between pre-resistant and pre-sensitive groups in each sample. To minimize experimental noise from cases where both sisters were sampled in the scRNA-seq analysis before the treatment, we performed the differential analysis only on the cells lacking sisters in the same pre-treatment samples.

### Pathway analysis

Gene expression signature gene lists were first ranked by adjusted p-value, then by non-adjusted p-value, and finally by log2FC. The ranked gene lists were subjected to Gene Set Enrichment Analysis (GSEA) using the classic enrichment statistic implemented in GSEA software (v4.3.2, University of California San Diego and Broad Institute) to inspect the MSigDB (Subramanian et al. 2005, Liberzon et al. 2011) v2023.2.Hs KEGG_LEGACY gene sets.

### Predicting drugs to target primed resistance

Four variants of pre-resistant gene expression signatures for both treatments were generated, including: 1) pre-resistance signature log2FC, 2) pre-resistance signature log2FC (gene list filtered by the Wilcoxon rank sum test *P*-value <0.05), 3) contrast log2FC between treatment and non-treatment control (pre-resistance signature log2FC - pre-fitness signature log2FC), and 4) contrast log2FC between treatment and non-treatment control (gene list filtered by bootstrapping test *P*-value < 0.05). Only sister-concordant genes were included in the analysis. Connectivity score defined in the Connectivity Map project (Lamb et al. 2006) was used to assess the similarity of these signatures and the drug-induced consensus gene expression signatures, constructed using LINCS L1000 Connectivity Map data, following the procedures described in previous work (Szalai et al. 2019, Douglass et al. 2022). Function “connectivityScore” (method = “fgsea”, nperm = 1000) from PharmacoGx R package (v3.0.2) (Smirnov et al. 2016) was used to calculate the connectivity scores. Drugs were ranked by their connectivity scores and filtered by *P*-value threshold <0.05 in all four pre-resistance signature variants.

### Drug combination testing and synergy scoring

Cell lines (600 or 800 cells/well for MOLM-13 and MV4-11, respectively) and primary patient samples (7500 cells/well) were seeded in 384-well plates in 25 µl of respective culture media using BioTek MultiFlo FX RAD (5 µl cassette) (Biotek, Winooski, VT, USA). Indicated concentrations of small molecules were added for 24 hours, after which indicated concentrations of midostaurin or quizartinib were added and incubated for 72 hours. Concentration ranges for drugs were chosen to induce 10% to 50% inhibition as single agents. As positive (total killing) and negative (growth control) controls 100 µM benzethonium chloride (MP Biomedicals, Santa Ana, CA, USA) and 0.2% DMSO were used, respectively. Cell viability was then determined using CellTiter-Glo 2.0 assay (Promega, Madison, WI, USA) according to manufacturer’s protocol. Each concentration and combination was tested in triplicate, and experiments were performed at least three times for cell lines and once or twice for patient samples. Representative results are shown.

Dose-response matrices were analyzed with SynergyFinder+ (https://tangsoftwarelab.shinyapps.io/synergyfinder/) (R package v3.8.2) (Zheng et al. 2022) using four different reference models, HSA (Highest Single Agent), Bliss (Bliss independence), Loewe (Loewe additivity), and ZIP (Zero Interaction Potency) to evaluate synergy between the drugs. HSA synergy scores were visualized as 3-D landscapes over the dose matrices. Mean synergy scores ≥ 5 were considered as synergistic. Significance of the mean synergy scores was assessed by bootstrapping (10 iterations).

### Western blotting

Cells were harvested by centrifugation, washed twice with cold PBS, and lysed with RIPA buffer (#89900, Thermo Fisher Scientific, Waltham, MA, USA) containing 2x Pierce protease inhibitor mixture (A32955, Thermo Fisher Scientific) and 1 mM EDTA on ice for 30 min. Lysates were sonicated and cleared by centrifugation. Protein concentration was determined with Pierce BCA protein assay kit (Thermo Fisher Scientific). Then, samples were boiled for 5 min in 1x Laemmli sample buffer and 5% 2-mercaptoethanol. Proteins were resolved on 4–20% Mini-PROTEAN® TGX™ precast protein gels (Bio-Rad) and transferred to Immobilon-FL PVDF membranes (Merck). After blocking with 2% BSA, membranes were incubated with the primary antibodies for 2h at RT or overnight at +4°C. Rabbit polyclonal antibody against GSPT1 (10763-1-AP, Proteintech), and as a loading controls mouse monoclonal antibodies against alpha-tubulin (DM1A, Abcam, Cambridge, UK) and alpha-actin (JLA20, Merck) were used as primary antibodies. Immunodetection was performed with Azure 500 imager (Azure Biosystems, Dublin, CA, USA) using IRDye secondary antibodies (LI-COR Biosciences, Lincoln, NE, USA). Protein band signals were quantified using ImageJ (1.53c, NIH, Bethesda, MD, USA).

## Acknowledgements

We thank prof. Caroline Heckman (University of Helsinki) for providing the MOLM-13 cell line. Authentication of the cell line was performed at Institute for Molecular Medicine Finland (FIMM) Genomics Unit of Technology Centre supported by Helsinki Institute of Life Science (HiLIFE) and Biocenter Finland. Recombinant virus work was performed at Biomedicum Virus Core (HelVi-BVC) supported by HiLIFE and the Faculty of Medicine, University of Helsinki, and Biocenter Finland. Single-cell transcriptomics was performed at FIMM Single-Cell Analytics unit supported by HiLIFE and Biocenter Finland. Figs. 1A, 4M, and 5A were created with BioRender.com. ChatGPT (OpenAI) was used with careful consideration to improve language and readability of the manuscript text. This work was supported by The Sigrid Jusélius Foundation (A.V., J.T.), The Cancer Foundation Finland (A.V., S.Z.), Foundation for the FinnishCancer Institute (K. Albin Johansson Cancer Research Fellowship for A.V.), Research Council of Finland projects 289059 (A.V.), 319243 (A.V.), 317680 (J.T.), 320131 (J.T.), 351165 and 351196 (J.T., A.V.), ERA PerMed JTC2020 PARIS/Research Council of Finland project 344697 (A.V.), University of Helsinki Research Foundation (J.D., S.Z.), K. Albin Johanssons stiftelse (S.Z.), Ida Montinin Säätiö (S.Z.), Biomedicum Helsinki Foundation (S.Z.). This study was co-funded by the European Union (J.T.: European Research Council project DrugComb No. 716063; AV: European Research Council project STRONGER, No. 101125261). Views and opinions expressed are however those of the author(s) only and do not necessarily reflect those of the European Union or the European Research Council. Neither the European Union nor the granting authority can be held responsible for them.

## Author contributions

**J. Eriksson**: Conceptualization, methodology, investigation, formal analysis, visualization, writing - original draft, writing - review and editing. **S. Zheng**: Formal analysis, visualization, writing - original draft, writing - review and editing. **J. Bao**: Investigation, writing - review and editing. **J. Dai**: Methodology, investigation, writing - review and editing. **W. Wang**: Data curation, writing - original draft, writing - review and editing. **A. Vähärautio**: Supervision, resources, writing - review and editing: **J. Tang**: Conceptualization, methodology, supervision, resources, writing - review and editing.

---

**Fig. S1.**
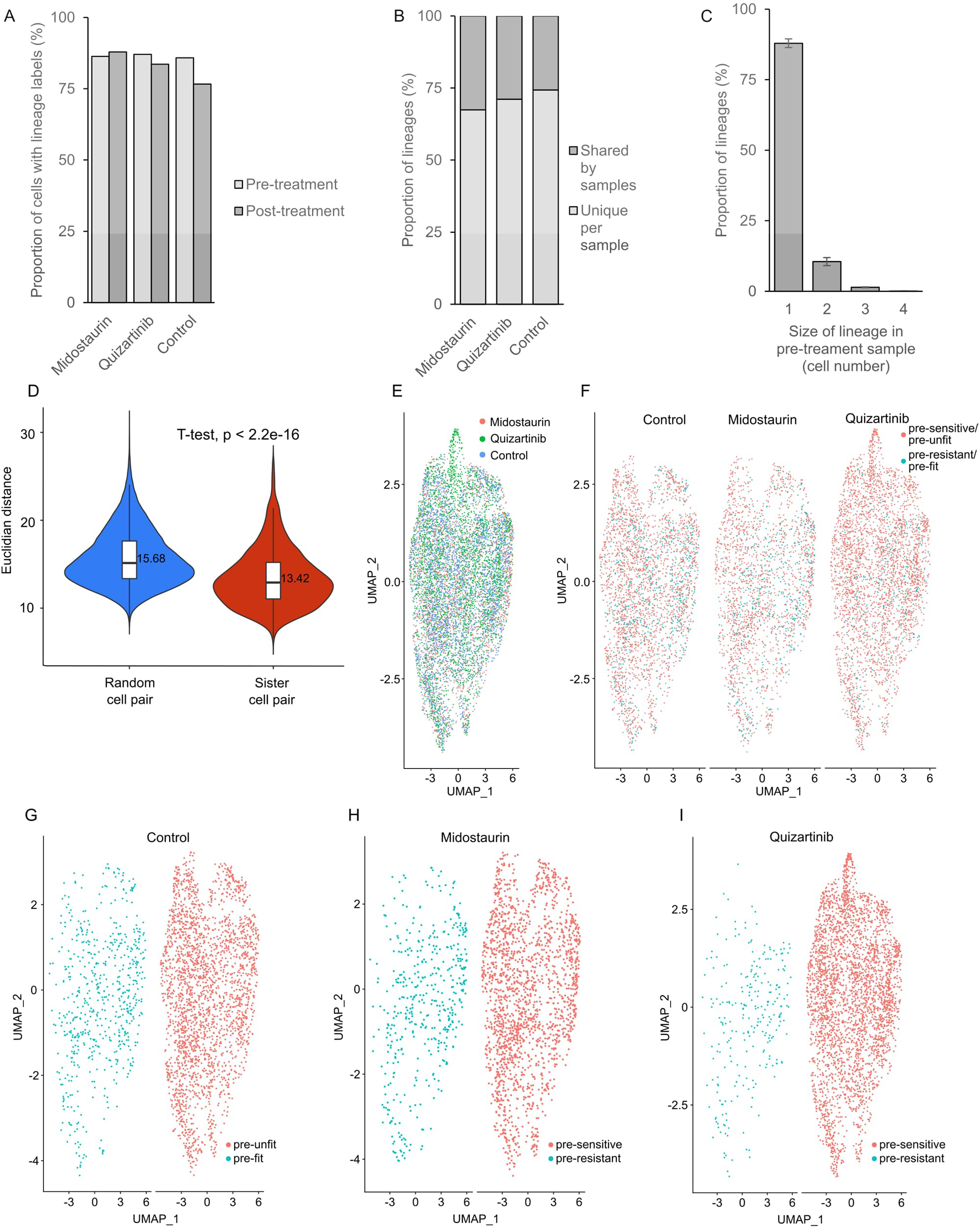
Lineage labeling efficiency, label detection assessment, and sister cell similarity. (A) Proportion of cells with lineage labels detected per sample. (B) Proportion of unique and shared barcodes in each pre-treatment sample. (C) Average proportion of lineages with indicated numbers of cells within a pre-treatment sample. (D) Euclidean distance of sister cell pairs in comparison to random cell pairs in the pre-treatment samples (*P* < 2.2e-16, two-tailed t-test). (E) UMAP projection of the pre-treatment samples. (F-G) UMAP projections of pre-resistant and pre-sensitive in each pre-treatment sample or pre-fit and pre-unfit cells in the DMSO-treated control.

**Fig. S2.**
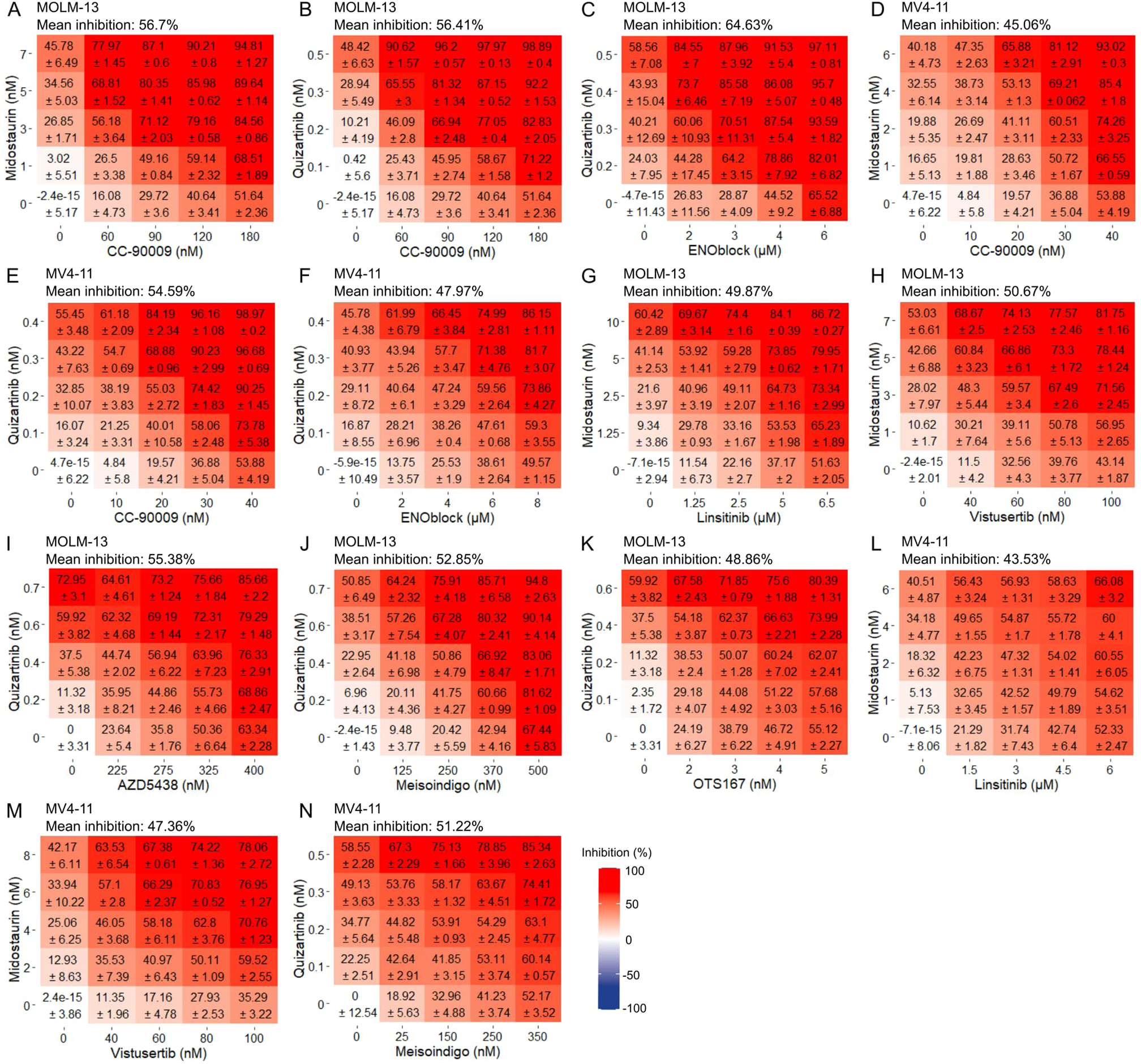
Dose-response matrices for drug combinations in MOLM-13 and MV4-11 cell lines. Dose-responses (mean ± standard deviation) for indicated drugs in combination with midostaurin or quizartinib in MOLM-13 (A-C, G-K) and MV4-11 (D-F, L-N) cells. Cells were first treated with the drug on horizontal axis with indicated concentrations, and after 24 hours midostaurin or quizartinib were added for 72 hours with concentrations indicated in the vertical axis.

**Fig. S3.**
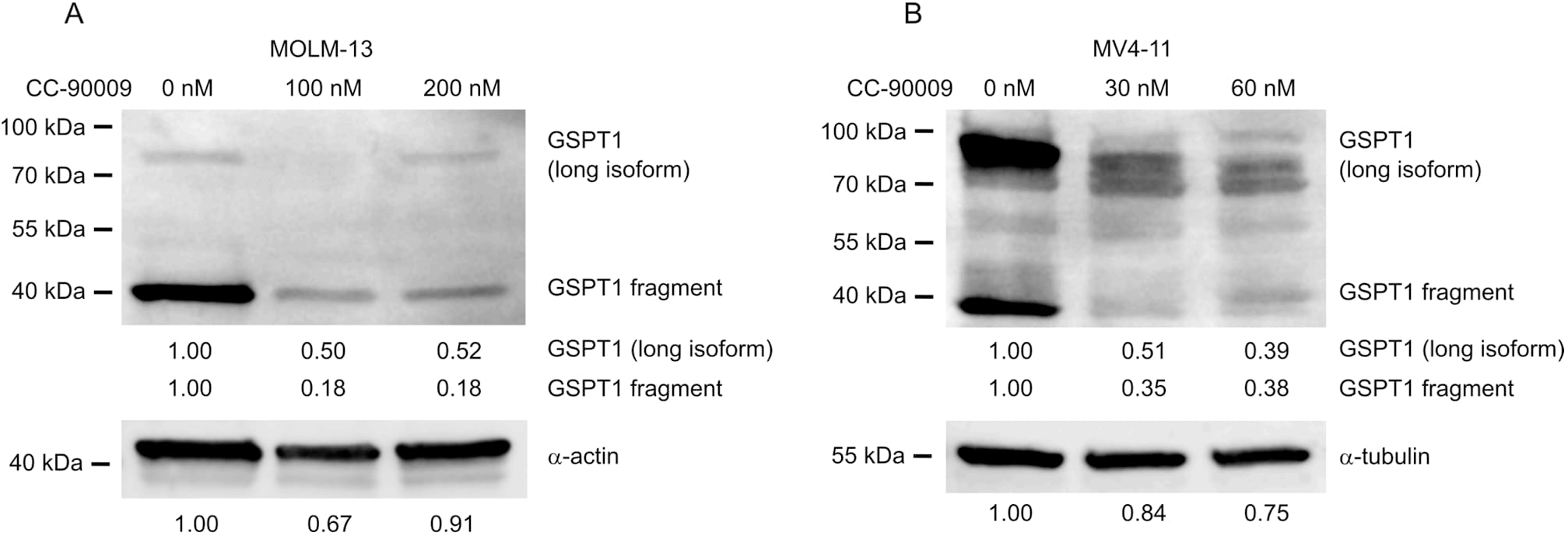
GSPT1 protein levels after CC-90009 treatment. Western blot analysis of GSPT1 protein levels in MOLM-13 (A) and MV4-11 (B) cell lines after treating the cells with DMSO or indicated concentrations of CC-90009 for 24 hours. Alpha-actin or alpha-tubulin were used as loading controls. Relative levels of each protein or protein fragment compared to the DMSO control are shown below.

**Fig. S4.**
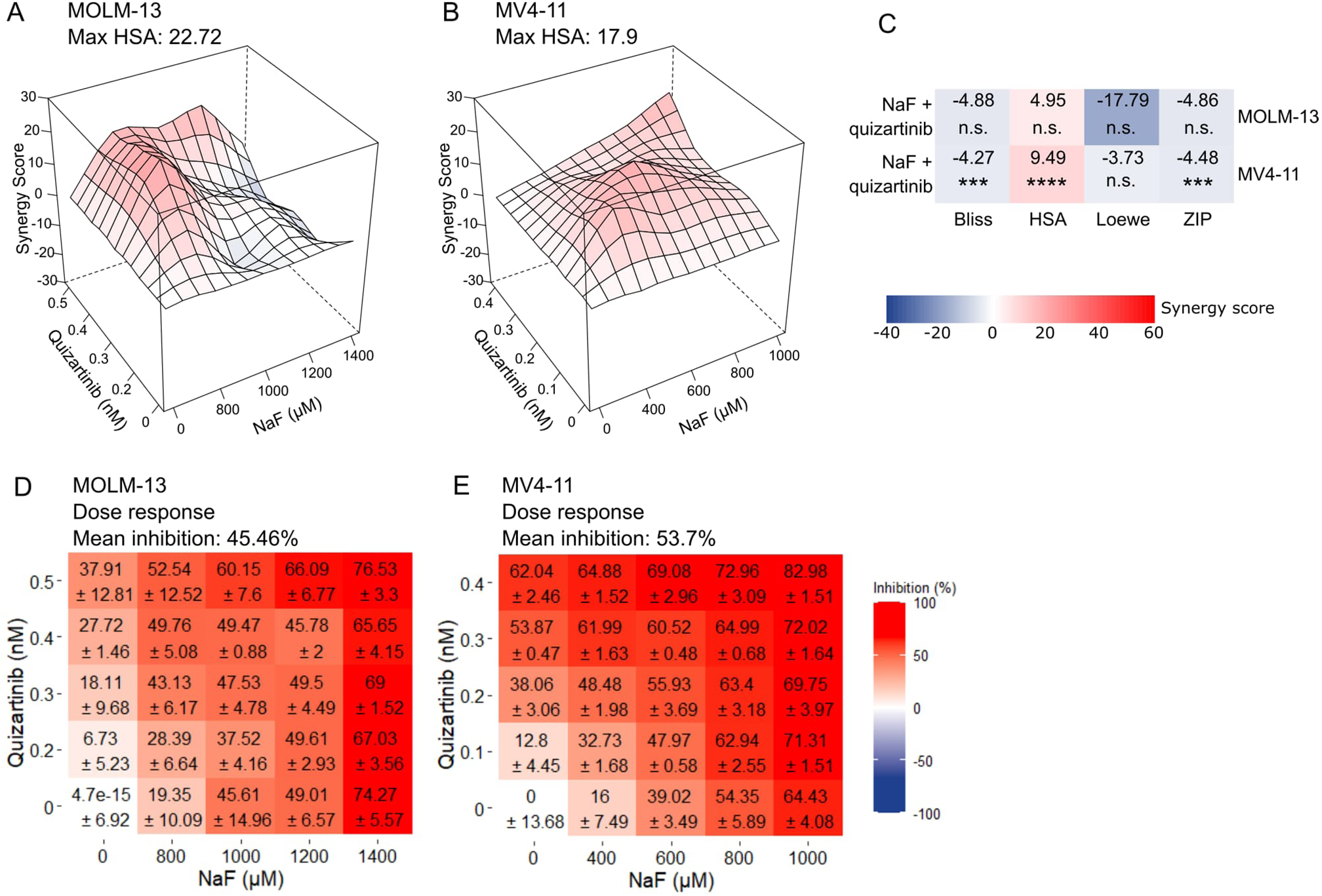
Evaluation of the synergism between sodium fluoride and quizartinib treatments. HSA synergy landscapes (A-B), mean synergy scores (C), and dose-response matrices (D-E) for sodium fluoride (NaF) in combination with quizartinib in MOLM-13 (A, C, D) and MV4-11 cells (B, C, E). (C) *P-*values were determined by bootstrapping test; n.s. = not significant, \**P* < 0.05, \*\**P* < 0.01, \*\*\**P* < 0.001, \*\*\*\**P* < 0.0001. (D-E) Dose-response matrices show mean inhibition (%) ± standard deviation.

**Fig. S5.**
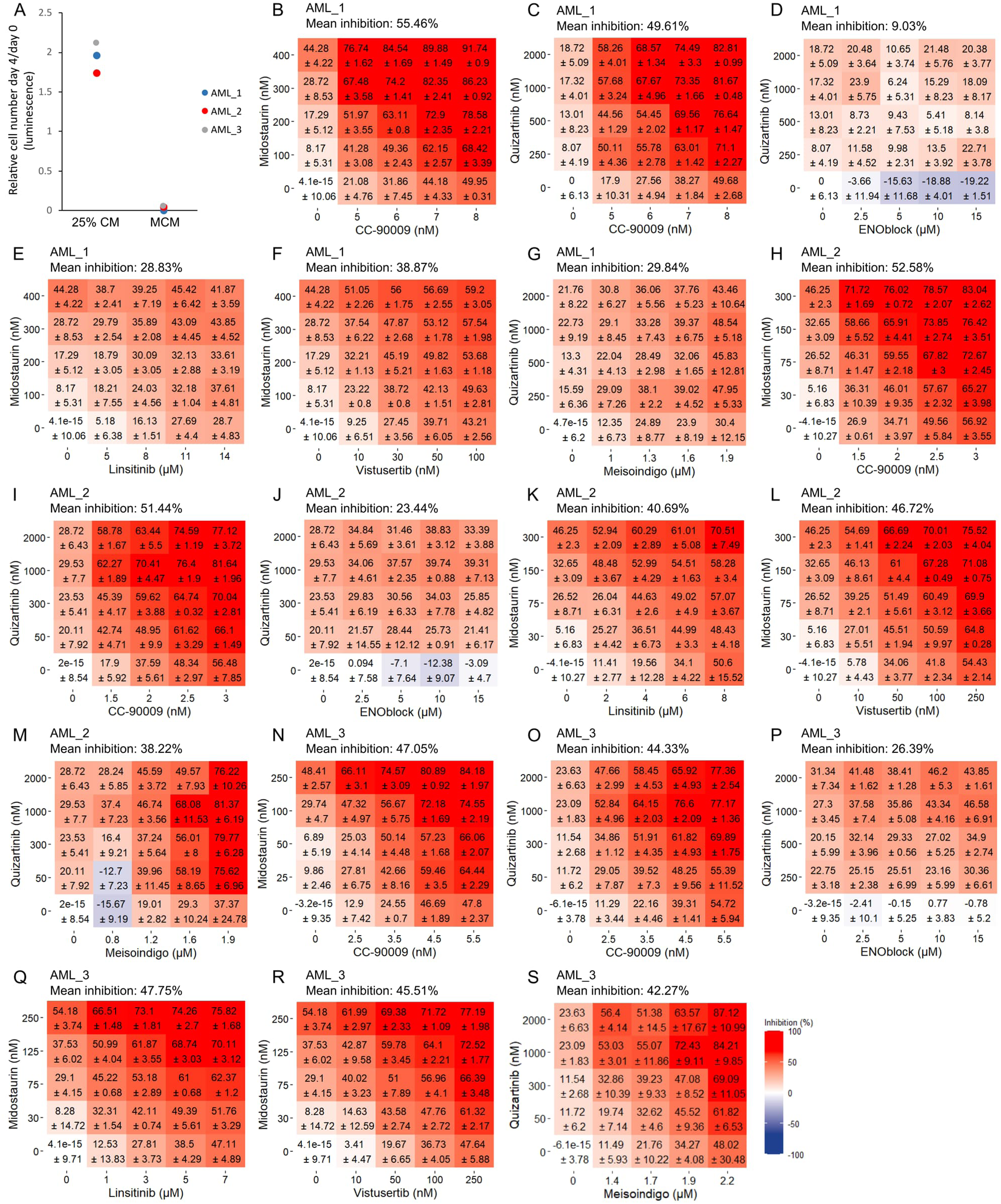
Drug combination testing in primary AML patient samples. (A) Relative cell number (CTG luminescence signal) in three FLT3-ITD-mutated AML patient samples after 4 days of culture in 25% HS-5 cell-conditioned medium (CM) or Mononuclear Cell Medium (MCM). (B-S) Dose-responses (mean inhibition % ± standard deviation) for indicated drugs in combination with midostaurin or quizartinib in patient sample AML_1 (B-G), AML_2 (H-M), and AML_3 (N-S). Cells were first treated with the drug on horizontal axis with indicated concentrations, and after 24 hours midostaurin or quizartinib were added for 72 hours with concentrations indicated in the vertical axis.

**Fig. S6.**
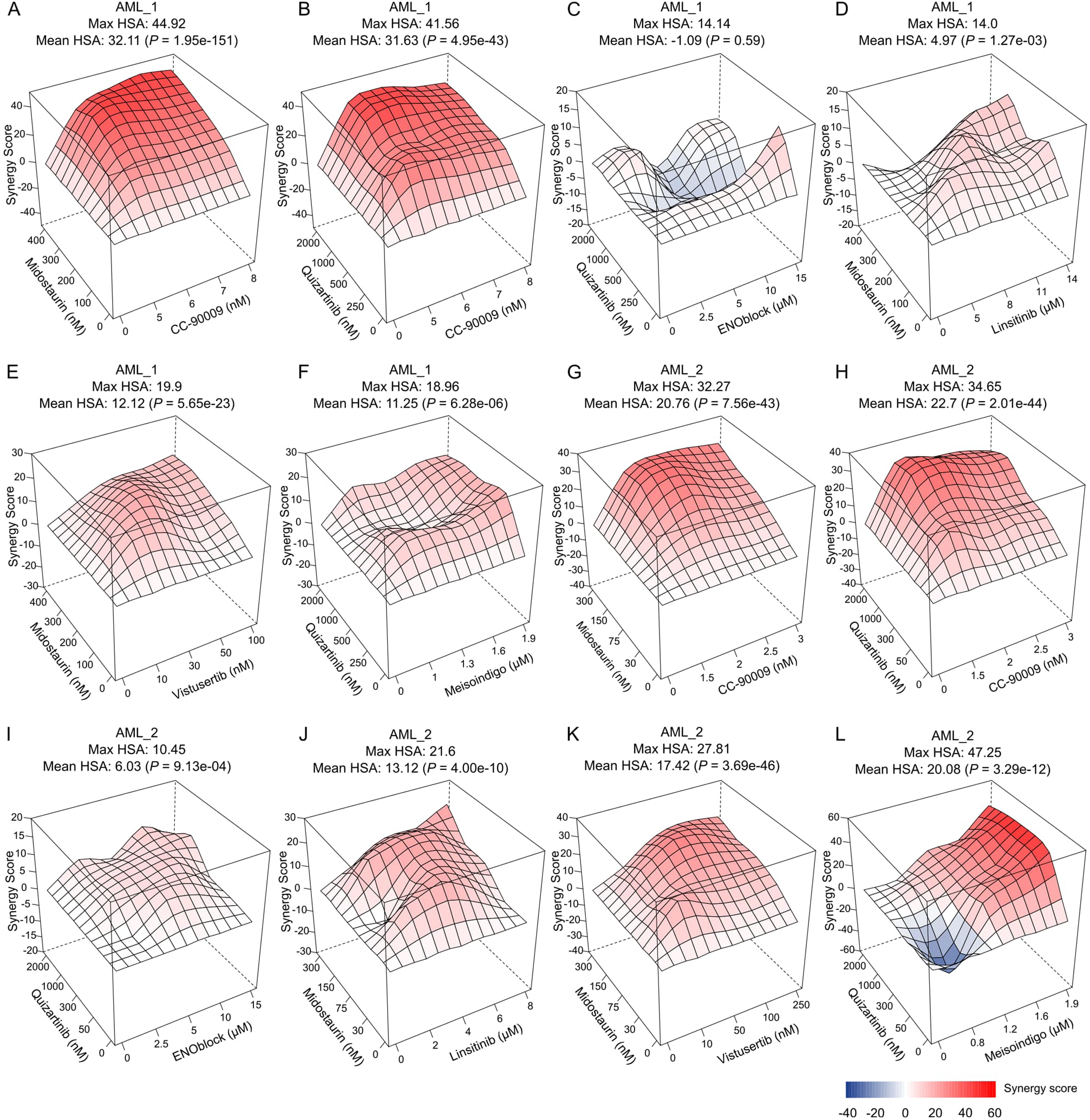
Synergistic effects of drug combinations in primary AML patient samples. HSA synergy landscapes for the indicated drugs in combination with midostaurin or quizartinib in primary AML patient samples AML_1 (A-F) and AML_2 (G-L). *P*-values were determined by bootstrapping test.

## Supplementary Tables

Table S1. Pre-resistance gene expression signatures for midostaurin treatment.

Table S2. Pre-resistance gene expression signatures for quizartinib treatment.

Table S3. Pre-resistance gene expression signatures for control condition.

Table S4. Full list of significant KEGG pathways for pre-resistance and pre-fitness (control) signatures.

Table S5. Full list of significant BioCarta pathways for pre-resistance and pre-fitness (control) signatures.

Table S6. Genes significantly and specifically up-regulated in midostaurin pre-resistant cells compared to both pre-sensitive cells and pre-fit signal within the DMSO control.

Table S7. Genes significantly and specifically up-regulated in quizartinib pre-resistant cells compared to both pre-sensitive cells and pre-fit signal within the DMSO control.

Table S8. Drugs predicted to sensitize to midostaurin and quizartinib treatments.

Table S9. Patient characteristics.

